# Haplotype divergence supports ancient asexuality in the oribatid mite *Oppiella nova*

**DOI:** 10.1101/2020.12.07.414623

**Authors:** A. Brandt, P. Tran Van, C. Bluhm, Y. Anselmetti, Z. Dumas, E. Figuet, C. M. François, N. Galtier, B. Heimburger, K. S. Jaron, M. Labédan, M. Maraun, D. J. Parker, M. Robinson-Rechavi, I. Schaefer, P. Simion, S. Scheu, T. Schwander, J. Bast, 2020

**Author notes:** Shared senior authorship, authors four to 16 in alphabetical order (S. Scheu, T. Schwander, J. Bast).

## Abstract

Sex strongly impacts genome evolution via recombination and segregation. In the absence of these processes, haplotypes within lineages of diploid organisms are predicted to accumulate mutations independently of each other and diverge over time. This so-called ‘Meselson effect’ is regarded as a strong indicator of the long-term evolution under obligate asexuality. Here, we present genomic and transcriptomic data of three populations of the asexual oribatid mite species *Oppiella nova* and its sexual relative *Oppiella subpectinata*. We document strikingly different patterns of haplotype divergence between the two species, strongly supporting Meselson effect like evolution and ancient asexuality in *O. nova*: (I) Variation within individuals exceeds variation between populations in *O. nova* but *vice versa* in *O. subpectinata*. (II) Two *O. nova* sub-lineages feature a high proportion of heterozygous genotypes and lineage-specific haplotypes, indicating that haplotypes diverged independently within the two lineages after their split. (III) The deepest split in gene trees generally separates haplotypes in *O. nova*, but populations in *O. subpectinata*. (IV) Tree topologies of the two haplotypes match each other. Our findings provide positive evidence for the absence of sex over evolutionary time in *O. nova* and suggest that asexual oribatid mites can escape the dead-end fate usually associated with asexual lineages.

## Introduction

Sexual reproduction is considered as a prerequisite for the long-term persistence of eukaryote species, because it reduces selective interference among loci and thus facilitates adaptation and purifying selection (Hill & Robertson, 1966; Felsenstein, 1974; Bell, 1982; Rice & Friberg, 2009; Neiman *et al*., 2017). Contrary to this scientific consensus, some exceptional taxa appear to have persisted in the absence of sex over millions of years, the so-called ‘ancient asexual scandals’ (sensu Judson & Normark, 1996; Schoen *et al*., 2009a; Schurko *et al*., 2009). These exceptional taxa are invaluable, because by understanding how they persisted as asexuals they could help to identify the adaptive value of sex (Butlin, 2002), one of the major riddles in evolutionary biology (Maynard Smith, 1978; Bell, 1982). However, several species originally believed to be ancient asexual scandals were later suggested to be either recently derived asexuals or to engage in some form of rare or non-canonical sex (Lunt, 2008; Schurko *et al*., 2009; Signorovitch *et al*., 2015; Schwander, 2016; Laine *et al*., 2020). At least two candidates for ancient asexuality remain, the darwinulid ostracods and several parthenogenetic lineages of oribatid mites. Both groups appear to have persisted for tens of millions of years (Heethoff *et al*., 2009; Schoen *et al*., 2009b) and diversified into ecologically different species (Birky & Barraclough, 2009; Schurko *et al*., 2009). However, support for obligate asexuality in darwinulid ostracods and oribatid mites has largely been based on negative evidence, i.e. the absence of males among thousands of females and the non-functionality of rare males (Taberly, 1988; Palmer & Norton, 1991; Birky, 2010; Wehner *et al*., 2018). Screening these groups for positive evidence of ancient asexuality is therefore of major importance (Normark *et al*., 2003).

One of the strongest predictions for evolution without recombination and segregation is that the two haplotypes (each stemming from one homologous chromosome copy) within a diploid clonal lineage accumulate mutations independently of each other. Thus, after the loss of sex, the haplotype sequences diverge over time, and levels of intra-individual heterozygosity increase (**Figure 1**). This intra-individual haplotype divergence is commonly known as the ‘Meselson effect’ (Birky, 1996; Normark *et al*., 2003).

**Figure 1:**
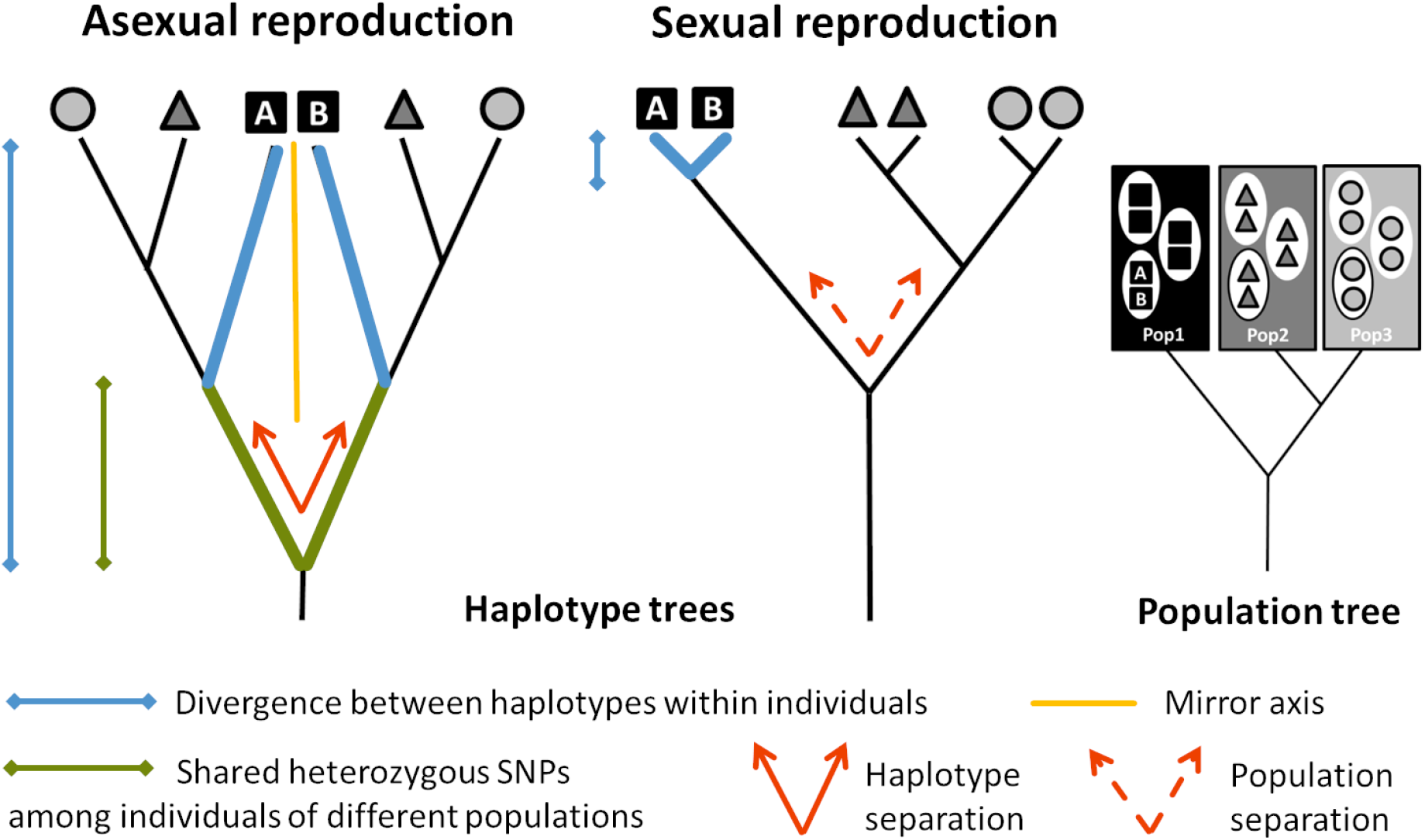
Nuclear haplotype trees expected under (long-term) obligate asexual and sexual reproduction. In diploid, functionally clonal organisms, homologous chromosomes accumulate mutations independently of each other and evolve as independent lineages (note that this can be restricted to specific regions sheltered from a loss of heterozygosity caused by mechanisms such as gene conversion). Accordingly, divergence between haplotypes within individuals (blue) is expected to exceed the mean divergence between haplotypes of individuals from different populations. Furthermore, the haplotype tree fully separates homologous haplotypes at its deepest split (red), which results in high frequency of heterozygous SNPs shared among individuals of different populations (green). Finally, the topologies of haplotype subtrees A & B are expected to match each other (the orange line represents the mirror axis) due to their parallel divergence. In sexual organisms, haplotype divergence is expected to follow population divergence and the haplotype tree to resemble that of the populations. Therefore, in sexuals, divergence between haplotypes within individuals is expected to be smaller than the divergence between populations, and the haplotype tree fully separates populations (red dashed). Figure adapted from (Schwander *et al*., 2011).

Surprisingly, there has only been equivocal empirical validation of this strong theoretical prediction thus far. In several asexual lineages the Meselson effect was not found (e.g., darwinulid ostracods; Schoen & Martens, 2003), or could be explained by mechanisms other than haplotype divergence after the transition to asexuality, such as a hybrid origin (e.g., *Meloidogyne* nematodes; Lunt, 2008) or divergence between paralogs (ohnologs) rather than between haplotypes (e.g., bdelloid rotifers; Mark Welch *et al*., 2008; Flot *et al*., 2013, and *Timema* stick insects; Schwander *et al*., 2011; Jaron *et al*., 2020a; reviewed in Hoerandl *et al*., 2020). Potential support for the Meselson effect was found in fissiparous species of *Dugesia* flatworms, but with data on only two genes, alternative explanations such as divergent paralogs could not be excluded (Leria *et al*., 2019). The as yet strongest support comes from a whole-genome study of obligately asexual trypanosomes, unicellular parasitic flagellates, in which some genomic regions are highly heterozygous and show the expected parallel haplotype divergence (Weir *et al*., 2016). We still lack any support from genome-wide analyses of the Meselson effect in asexual animals.

One of the most promising eukaryotic systems for understanding long-term persistence in the absence of sex are oribatid mites (Heethoff *et al*., 2009; Schwander, 2016). Oribatid mites are small (150-1400 μm), soil living chelicerates that play an important role as abundant decomposers in most terrestrial ecosystems (Heethoff *et al*., 2009; Maraun *et al*., 2012, 2019). A number of lineages lost sex independently, providing the possibility for comparative analyses (Norton & Palmer, 1991; Cianciolo & Norton, 2006; Pachl *et al*., 2020). As yet, the cellular mechanism underlying asexuality in oribatid mites has not been determined with certainty. Cytological studies, focussed mostly on a single species (*Archegozetes longisetosus*), have suggested a modified meiosis (holocentric chromosomes undergoing terminal fusion automixis with an inverted sequence of meiotic divisions) that preserves heterozygosity in regions sheltered from recombination and other homogenising mechanisms (Taberly, 1987; Wrensch *et al*., 1993; Heethoff *et al*., 2006; Laumann *et al*., 2008; Engelstaedter, 2017; Bergmann *et al*., 2018).

In this study, we characterize haplotype divergence patterns in the asexual oribatid mite species *Oppiella nova* and its sexual relative *O. subpectinata*. A previous study, based on molecular divergence estimates, suggested that *O. nova* persisted in the absence of sex for millions of years, given that sub-lineages within this species split 6 to 16 myr ago, (Schaefer *et al*., 2010; Von Saltzwedel *et al*., 2014). Using *de novo* genomes and polymorphism data from transcriptome-resequencing, we tested for four population genomic signatures expected under obligate asexuality (Meselson effect; see **Figure 1**). These signatures should be present in the asexual species *O. nova*, but absent in its close sexual relative *O. subpectinata*: (I) high divergence within individuals, exceeding the divergence between populations (**Figure 1**; blue); (II) high frequency of shared heterozygous variants among individuals of different populations (indicating that haplotypes diverged prior to separation of populations; **Figure 1**; green); (III) the deepest split in allele phylogenies separates haplotypes, not populations (as opposed to sexual organisms where the deepest split typically separates populations; **Figure 1**; red); (IV) the topologies of trees based on haplotypes A & B match each other due to parallel divergence of haplotypes during population separation (**Figure 1**; orange).

## Results

### De novo genomes

We first *de novo* assembled genomes of single individuals of the asexual oribatid mite species *O. nova* and of its sexual relative *O. subpectinata*. The quality- and contaminant-filtered assemblies (v03, see **Data availability**) spanned a total size of 197 Mb for *O. nova* and 213 Mb for *O. subpectinata*. They contained 63,118 and 60,250 scaffolds, with an N_50_ of 6,753 and 7,017 bp, with 23,761 and 23,555 genes annotated(see **Supplementary Table S1, S2, S3** and **Methods** for details). Despite the low assembly contiguity (likely caused by whole genome amplification), the genomes were of sufficient quality for down-stream analyses which focused on patterns of heterozygosity and polymorphism in genes. This is reflected by the high completeness scores of arthropod core genes (C) as inferred via BUSCO, with few fragmented (F), missing (M) or duplicated (D) genes (C: 87.5%, F: 6.6%, M: 5.9%, D: 8.6% for *O. nova* and C: 86.2%, F: 6.4%, M: 7.4%, D: 7.9% for *O. subpectinata* see **Supplementary Table S1)**.

### (I) The divergence within asexual individuals is large and exceeds the divergence between populations

We analysed within-individual and between-population divergence of individuals from three geographically distant locations in Germany (Hainich, H; Kranichstein Forest, KF; Schwaebische Alb, SA), using transcriptome data from three individuals per species and location (for sampling sites see **Supplementary Figure 1** and **Methods**). Polymorphism data were generated by mapping transcriptome reads of each individual to the corresponding reference genome. For *O. nova*, a multi-dimensional scaling plot (MDS; based on raw Hamming distances) revealed the presence of at least two clusters (hereafter referred to as divergent lineages), grouping individuals from different sampling locations (**Figure 2a**). For *O. subpectinata*, individuals were separated into three distinct clusters, each corresponding to one location (**Figure 2b**). Accordingly, between-location variation contributed much less to the species-wide genetic variation in *O. nova* (12.0%) than in *O. subpectinata* (56.4%). In *O. nova* most variation (90.8%) was explained by variation within individuals.

**Figure 2:**
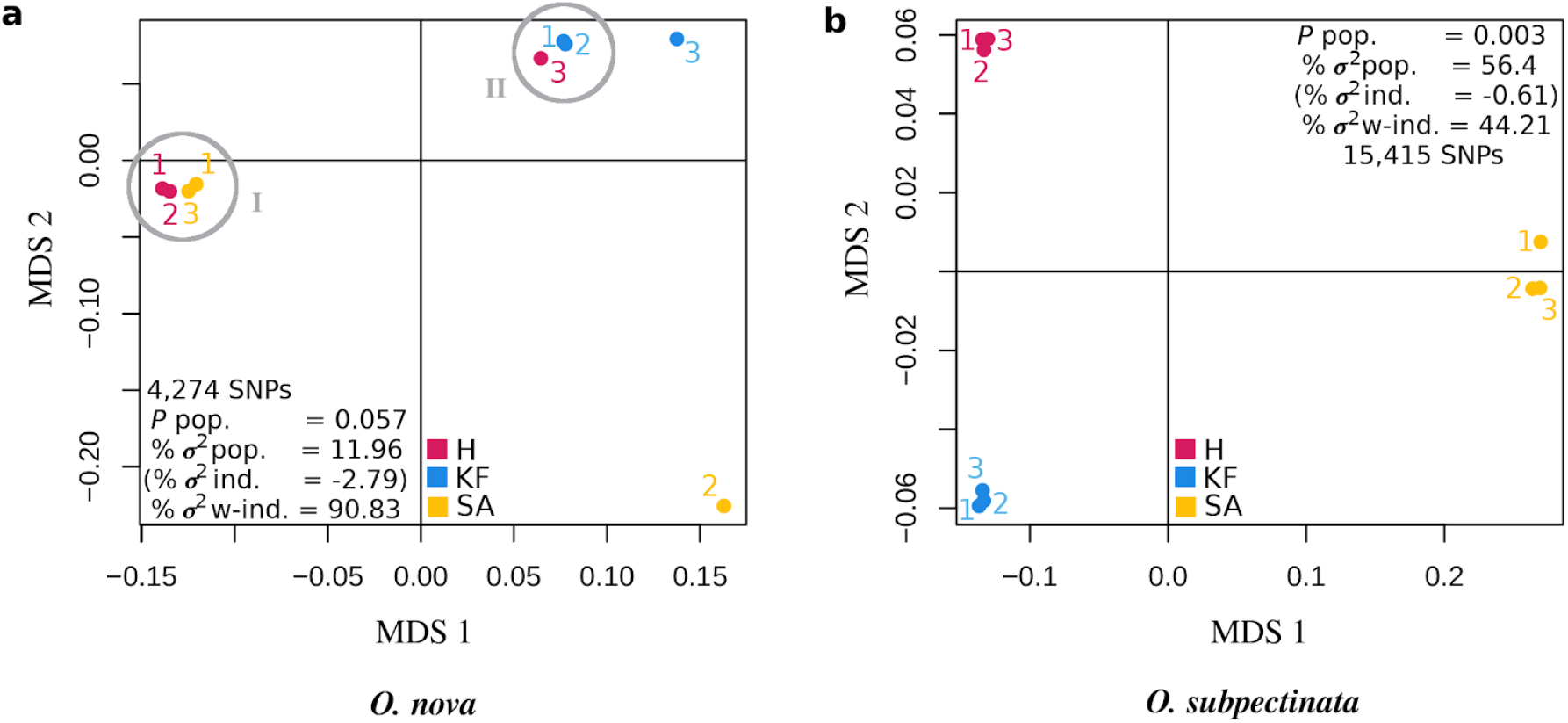
Genetic divergence is more extensive within individuals than between populations for the asexual *Oppiella nova* **(a)**, in contrast to the sexual *Oppiella subpectinata* **(b)**. In *O. nova* there are multiple genetic lineages grouping individuals from different geographical locations. Lineages are represented by two clusters and two single individuals (lineages one and two highlighted by grey circles; non-significant between-population variation; rand-test *P* pop. = 0.057). Two *O. nova* individuals, individual 3 from location KF and individual 2 from location SA, are rather homozygous and likely do not feature the Meselson effect, while the remaining individuals do (see Result sections II - IV). Individuals of the sexual *O. subpectinata* clustered by location (significant between-population variation; rand-test *P* pop. = 0.003). Notably, the majority of total genetic variation is explained by differences between populations (% ***σ***^2^ pop.) in *O. subpectinata*, but by within-individual differences in *O. nova* (% ***σ***^2^ w-ind.; % ***σ***^2^ ind.: % variation between individuals within location).

Consistent with the large proportion of intra-individual variation in *O. nova*, heterozygosity for most (seven out of nine) *O. nova* individuals was higher than heterozygosity for any of the nine *O. subpectinata* individuals (**Table 1**). This is the expected pattern given that haplotype divergence under functionally mitotic asexuality should result in increased heterozygosity levels compared to closely related sexual species, unless gene conversion or other homogenizing mechanisms occur at higher rates than new mutations (Birky, 1996).

**Table 1:** Individual heterozygosity estimates as percentages of heterozygous sites among all sites with available SNP genotypes for all nine individuals (inferred from transcriptome data; see also **Supplementary Figure 2**).

| location | *O. nova* (126,196 sites) |  | *O. subpectinata* (355,249 sites) |  |
| --- | --- | --- | --- | --- |
|  | individual | % heterozygous sites | individual | % heterozygous sites |
| Hainich | H1 | 1.273 | H1 | 0.619 |
|  | H2 | 1.285 | H2 | 0.665 |
|  | H3 | 0.741 | H3 | 0.640 |
| Kranichstein forest | KF1 | 0.746 | KF1 | 0.599 |
|  | KF2 | 0.771 | KF2 | 0.581 |
|  | KF3 | 0.414 | KF3 | 0.591 |
| Schwaebische Alb | SA1 | 1.268 | SA1 | 0.723 |
|  | SA2 | 0.441 | SA2 | 0.685 |
|  | SA3 | 1.288 | SA3 | 0.743 |

### (II) An excess of shared heterozygous variants indicates that haplotypes diverged prior to lineages

The relation between individual heterozygosity and (sub-)population allele frequencies is expected to differ between obligate asexual organisms and panmictic sexual populations, with higher individual heterozygosity relative to population allele frequencies in asexuals (Balloux *et al*., 2003; Schurko *et al*., 2009). We compared observed individual heterozygosity *vs* heterozygosity expected from allele frequencies using F_is_ as a measure. We based estimates of expected heterozygosity on genetically differentiated locations in *O. subpectinata* but on genetically differentiated lineages (I + II) in *O. nova*, because genetic differentiation between locations was low in *O. nova* (see **Figure 2a**). F_is_ values were strongly negative for *O. nova* individuals (mean F_is_ = −0.328, **Table 2**). Such negative F_is_ values indicate an excess of observed heterozygosity, as expected for functionally clonal organisms. For *O. subpectinata*, values were in the range expected for a non-inbred panmictic sexual species (mean F_is_ = 0.002, **Table 2**) and values for *O. nova* were significantly lower (mean F_is_ = −0.328; Wilcox rank sum test, *W* = 0; *P* < 0.001).

**Table 2:** Per individual inbreeding coefficient estimates (F_is_). Estimates of F_is_ were based on location in *Oppiella subpectinata*, but on genetically divergent lineages in *Oppiella nova* (lineages I + II; lineage II bold + italicised). Note that it was not possible to estimate F_is_ for *O. nova* KF3 and SA2 because they likely represent divergent lineages on their own, and estimating F_is_ requires a (sub-)population context.

| location | *O. nova* |  | *O. subpectinata* |  |
| --- | --- | --- | --- | --- |
|  | *individual* | *Fis* (# sites) | *individual* | *Fis* (# sites) |
| Hainich | H1 | -0.390 (5,414) | H1 | 0.015 (12,730) |
|  | H2 | -0.361 (5,414) | H2 | -0.034 (12,730) |
|  | H3 | **-0.243 (3471)** | H3 | 0.001 (12,730) |
| Kranichstein forest | KF1 | **-0.246 (3471)** | KF1 | -0.006 (40,145) |
|  | KF2 | **-0.296 (3471)** | KF2 | 0.023 (40,145) |
|  | KF3 | NA | KF3 | 0.002 (40,145) |
| Schwaebisch Alb | SA1 | -0.361 (5,414) | SA1 | -0.018 (10,197) |
|  | SA2 | NA | SA2 | 0.058 (10,197) |
|  | SA3 | -0.398 (5,414) | SA3 | -0.019 (10,197) |

The extensive heterozygosity variation among *O. nova* individuals suggests that within a single origin of asexuality, the evolution of heterozygosity can follow strikingly different trajectories in different lineages. Independently of the potential causes driving the heterozygosity loss in some *O. nova* lineages (represented by individuals KF3, SA2), it is important to note that highly homozygous lineages cannot feature the Meselson effect as the rate of heterozygosity loss is greater or equivalent to the gain of heterozygosity via new mutations. We therefore conducted explicit tests for the Meselson effect solely on the seven of the nine individuals where it could potentially be present.

If the loss of sex in *O. nova* occurred prior to population separation, the observed heterozygosity excess in seven individuals is expected to result from shared heterozygous SNPs among individuals of the two different lineages (see green lines in **Figure 1**). To test this, we generated a Site Frequency Spectrum (SFS). Sites with heterozygous SNPs shared among the seven individuals were 19 times more frequent than expected under Hardy Weinberg Equilibrium (HWE; yellow bar in **Figure 3**). Furthermore, there was an excess of sites with heterozygous SNPs exclusively shared among the four individuals of lineage I (48 and eight times more frequent than expected under HWE; turquoise bars in **Figure 3**) or among the three individuals of lineage II (eleven and 35 times more frequent than expected under HWE; purple bars in **Figure 3**). These results are consistent with the accumulation of heterozygous variants after the loss of sex, followed by lineage divergence and independent accumulation of heterozygous variants within lineages I and II (see inset tree; **Figure 3**).

**Figure 3:**
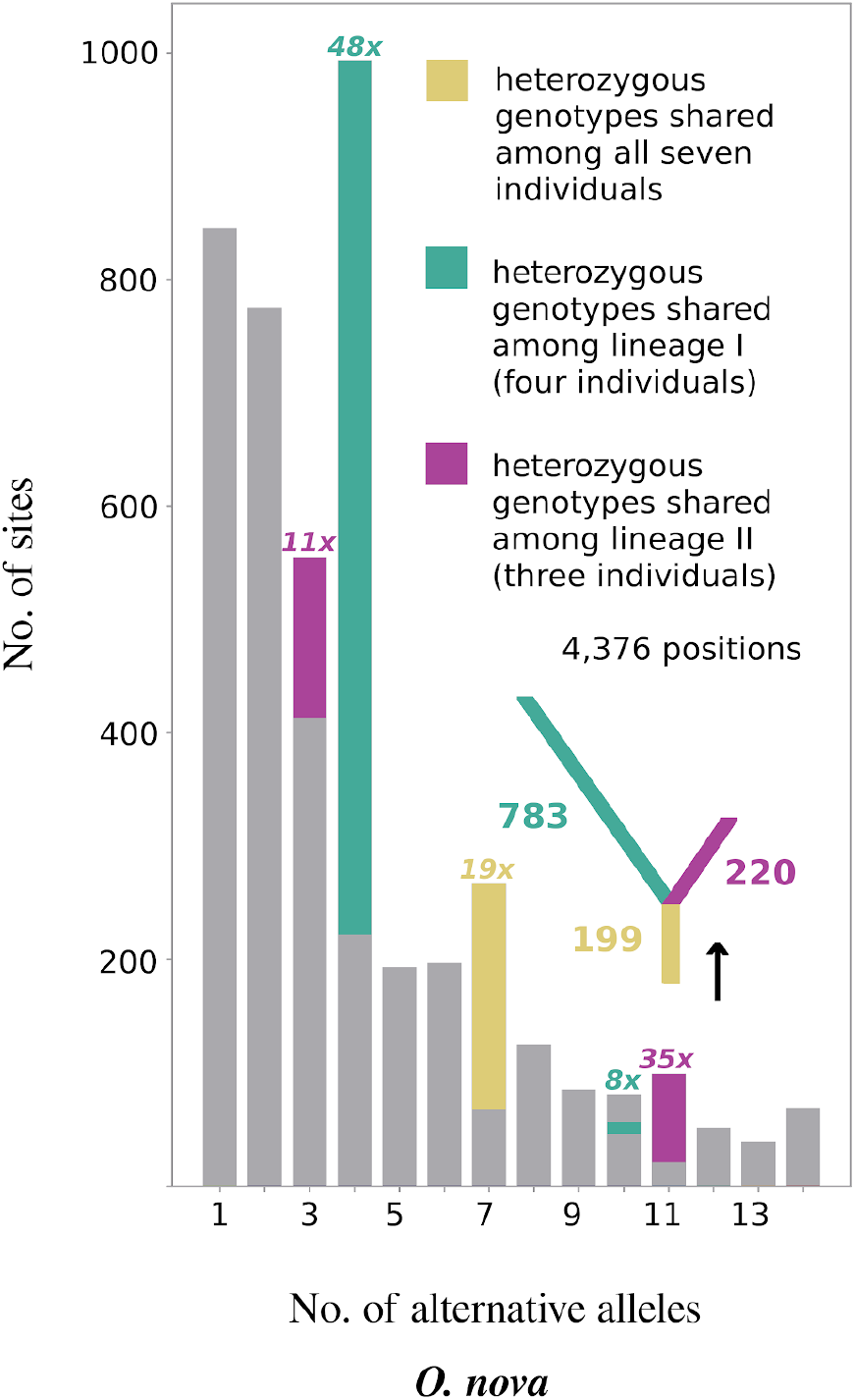
Excess of shared heterozyous SNPs among individuals of different populations and lineages for the asexual species *Oppiella nova*. The Site Frequency Spectrum (SFS) depicts the number of sites with a given number of non-reference variants over the seven heterozygous individuals (e.g., seven diploid individuals can display a maximum of 14 variants relative to the reference genome). Heterozygous genotypes shared among all seven individuals, or among individuals of lineages I and II privately, are colour-highlighted and their excess over HWE indicated (see legend). The SFS is consistent with the accumulation of shared heterozygous variants after the loss of sex, followed by lineage separation and independent accumulation of further heterozygosity in each lineage (inset tree with numbers of shared heterozygous variants at each branch).

### (III) The deepest split in many haplotype phylogenies separates haplotypes

A classical signature of haplotype divergence under asexuality is the full separation of haplotypes A and B at the deepest split of a haplotype tree. By contrast, haplotypes are generally expected to diverge according to population divergence in sexual organisms (**Figure 1**). To test these predictions, we phased haplotypes of the seven heterozygous individuals from the two genetic lineages of *O. nova* (which potentially show the Meselson effect based on % heterozygous sites and F_is_ estimates; see above) and the nine individuals of *O. subpectinata* using the RNAseq polymorphism data. We based all analyses on genomic regions with phases that formed a continuous overlap by at least 100 bp between at least two individuals (see **Methods**). For *O. nova* a total of 281 genome regions (median length 358 bp) were phased, spanning 140,966 bp and representing 86.3% of 163,418 theoretically phaseable sites (genotypes with coverage >= 10 for all seven individuals; for detailed information on phaseable regions see **Supplementary Table S4** and **Methods**). For *O. subpectinata*, a total of 275 regions (median length 563 bp) were phased. The regions spanned 206,255 bp, representing 58.1% of 355,249 theoretically phaseable sites, consistent with the considerably lower heterozygosity compared to *O. nova*.

To confirm that the phased regions represent allele-haplotypes and not merged paralogs, we verified that the genomic read coverage of phaseable regions was not exceeding the coverage of single-copy genes from the BUSCO arthropod database (merged paralogs should have an at least 2x higher coverage, see **Methods**). This was indeed the case as single-copy BUSCO genes, phased regions and the scaffolds from which phased regions derived showed a median coverage of 126x, 101x, and 87x, respectively. By contrast, the median coverage of known duplicate BUSCO genes, which served as a positive control, was about two-fold higher (207x) (**Supplementary Figure 3**).

We next aligned the phased haplotype sequences and calculated best fitting ML trees. We then computed for each tree how distinct it was from two topology-constrained trees (Kuhner & Felsenstein, 1994): (1) fully separating haplotypes A & B at the deepest split of the haplotype phylogeny, followed by population separation (consistent with predicted haplotype divergence under asexuality; hereafter referred to as “asex-tree”; **Figure 1)**, and (2) separating haplotypes according to population divergence as expected for a sexual organism (hereafter referred to as “sex-tree”; **Figure 4b**; **Figure 1**). We accounted for the observed coexistence of lineages I + II in *O. nova* by introducing lineage as an additional divergence level (**Figure 4a** blue trees; for *O. subpectinata* red trees). The delta of the two tree distance scores is indicative of a phaseable region being more consistent with haplotype divergence under asexuality (**Δ** _dist. asex-tree − dist. sex-tree_ < 0) or sexuality (**Δ** ._dista. sex-tree - dist. sex-tree_> 0). For *O. nova*, 115 regions (51.6%) showed a Delta < 0, while for the sexual *O. subpectinata*, this applied for only six regions (2.2%; see **Figure 4**). These results are corroborated by tree topology tests showing 69 of 223 phaseable regions being significantly more consistent with the asex-tree in *O. nova* but only one of 268 phaseable regions in *O. subpectinata* (for details see **Supplementary Table S5, Methods**). The 69 regions of *O. nova* spanned a total of 37,693 of the 163,418 theoretically phaseable sites, indicating that approximately 23.1% of the *O. nova* transcriptome shows a significantly better fit with the expected haplotype divergence pattern under asexuality than under sexual reproduction.

**Figure 4:**
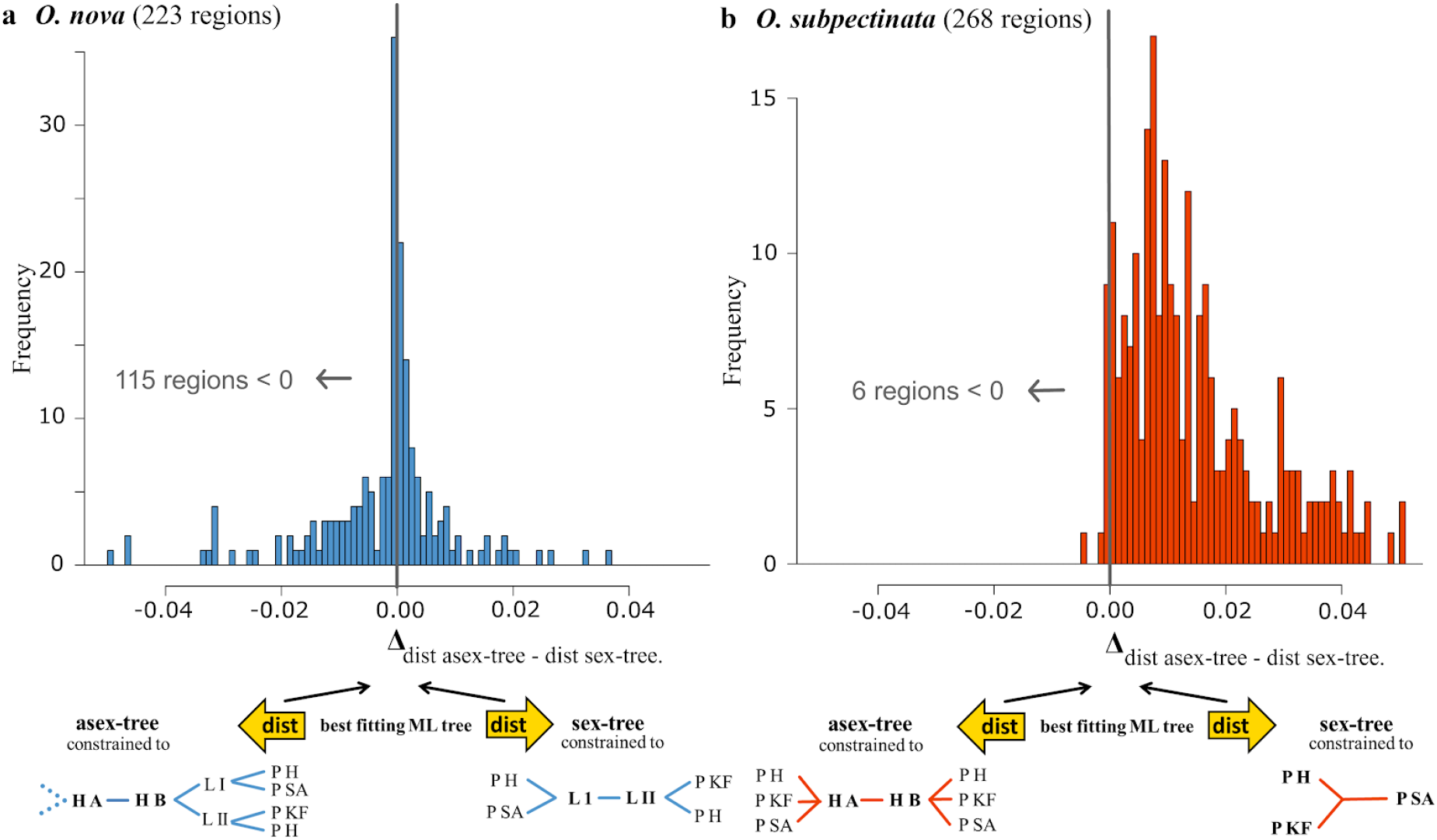
Haplotype trees are more consistent with asexuality in *Oppiella nova* **(a)** but with sex in *Oppiella subpectinata* **(b)**. Frequency distribution of per-region tree-distance score comparisons (Δ _dist. asex-tree - dist. sex-tree_). The score measures the combined distance (dist) in topology and branch lengths between an unconstrained tree and one of two constrained trees (asex-tree, sex-tree; see schematic trees for each species, respectively). A negative value indicates that a phaseable region’s best ML tree is more similar to its asex-tree than to its sex-tree. Reconstruction of constrained trees was possible for regions with >= 4 unique aligned sequences present, i.e. 223 and 268 regions for *O. nova* and *O. subpectinata*, respectively (**Methods**). To improve legibility, the histogram ranges are limited from −0.05 to 0.05, thereby excluding 26 regions below and eight regions above this range for *O. nova* and one region below and 32 regions above the range for *O. subpectinata*. H A: Haplotype A; H B: Haplotype B; L I: Lineage I; L II; Lineage II; P H; population Hainich; P KF; population Kranichstein forest; P SA; population Schwaebische Alb; dashed branches: lineage separation followed by population separation as for haplotype B.

### IV) The topologies of haplotype subtrees are matching

The high frequency of heterozygous genotypes shared among the seven individuals (**Figure 3**) and the split of haplotypes A and B at the basis of a haplotype tree (**Figure 4a**) could theoretically be explained by hybridisation at the origin of asexuality, while the separation of the two lineages I and II could have been caused by the loss of heterozygosity at different sites within each lineage. Therefore, we also tested for evidence of long-term asexual evolution within lineages by assessing parallel divergence of haplotypes within lineages I and II (comprising four individuals from populations H and SA and three individuals from populations H and KF). Thirty-one out of 39 resolved trees within lineage I fully separated haplotypes A & B, and 8 out of the 31 featured parallel divergence of haplotypes following the split. The 8 instances of parallel divergence are about 4 times more frequent than expected by chance, indicating that parallel divergence is a significant feature of lineage I (P < 0.001). For lineage II, 55 out of 90 resolved trees fully separated haplotypes, and 38 out of 55 featured parallel divergence. The 38 instances of parallel divergence are more than 2 times more frequent than expected by chance, meaning parallel divergence is also a significant feature of lineage II (P < 0.001).

## Discussion

Independent mutation accumulation in haplotypes of diploid asexual organisms is considered to be strong direct support for evolution under obligate asexuality (Birky, 1996; Normark *et al*., 2003). Surprisingly, empirical evidence for this ‘Meselson effect’ in parthenogenetic animals is, as yet, either lacking or equivocal (recently reviewed in Hoerandl *et al*., 2020). Here, we report population genomic signatures supporting the presence of the Meselson effect in the ancient asexual oribatid mite species *O. nova*, namely: (I) intra-individual variation exceeding between-population variation, (II) a large proportion of conserved heterozygous variants shared among individuals of different lineages and geographic locations, (III) separation of haplotypes rather than lineages in haplotype phylogenies, and (IV) topologies of haplotype subtrees are matching. These signatures were absent in the sexual species *O. subpectinata*. Accordingly, transcriptome-wide heterozygosity was overall higher in *O. nova* than in *O. subpectinata* even though two individuals of *O. nova* featured very low heterozygosity. To the best of our knowledge, this study is the first to provide strong positive evidence for the Meselson effect in a parthenogenetic animal and thus long-term evolution in the absence of sex.

Hybridization at the origin of asexuality can generate allele divergence patterns mimicking the Meselson effect even in recently evolved asexual species (Taylor & Foighil, 2000; Delmotte *et al*., 2003; Johnson, 2006; Lunt, 2008; Ament-Velásquez *et al*., 2016). However, a hybrid origin of asexuality is unlikely to explain our results for two reasons: first, asexual animals with a hybrid origin typically display within-individual divergence levels far exceeding the estimates in *O. nova* (Jaron *et al*., 2020b). Second, even if asexuality in *O. nova* was of hybrid origin, a hybrid origin accounts neither for the high frequencies of heterozygous SNPs private to the two divergent lineages (**Figure 3**) nor for the parallel divergence of haplotypes within lineages. Both patterns are likely generated by mutations that occurred after the origin of asexuality, which thus indicates long-term asexual evolution independently of possible hybridisation at the origin of asexuality.

Our results indicate that levels of haplotype divergence vary strongly among genomic regions in *O. nova* individuals, as well as among different *O. nova* lineages **(Figure 4a, Table 1**). This is in line with previous findings of varying levels of heterozygosity loss *vs* retention in other asexual animal species (Jaron *et al*., 2020b) and could explain why previous studies using individual genes in asexual mites and ostracods found no increased heterozygosity (Schoen & Martens, 2003; Schaefer *et al*., 2006). Our results thus illustrate that haplotype divergence and other genomic consequences of asexuality need to be studied on whole genomes or transcriptomes rather than on a few genes (see also Neiman *et al*., 2018).

Besides haplotype divergence in *O. nova*, our study indicates the presence of coexisting, strongly divergent lineages with different heterozygosity levels (**Figure 2a; Table 1**). Coexistence of strongly divergent lineages in *O. nova* has been shown previously based on the mitochondrial gene COI (separation of lineages was estimated to have occurred 6-16 mya ago) and was considered to indicate coexistence of forest and grassland genotypes (Von Saltzwedel *et al*., 2014). *O. nova* occurs over a wide variety of habitat types and shows a cosmopolitan distribution (Subias, 2004), suggesting that the extensive intra-specific polymorphism might be linked to large population sizes (Brandt *et al*., 2017). Independently of the origin of the extensive polymorphism, we also observed differences in heterozygosity between lineages. These could be linked to different mutation rates between lineages or to the presence of non-mutually exclusive mechanisms of heterozygosity loss, including the meiotic parthenogenesis proposed for asexual oribatid mites, lineage-specific deletions (hemizygous genome regions; Xu *et al*., 2010), and (GC-biased) gene conversion (Marais, 2003). GC-biased gene conversion has notably been suggested to contribute to the loss of heterozygosity in other parthenogenetic animals, e.g. in the darwinulid ostracod *Darwinula stevensoni* and in aphids of the tribe *Tramini* (Normark, 1999; Schoen *et al*., 2003).

Irrespective of the mechanisms underlying heterozygosity losses in *O. nova*, our results strongly support genome evolution in the absence of sex over evolutionary times in the asexual oribatid mite species *O. nova*. This is in line with previous studies which have shown that oribatid mites are able to overcome some major selective disadvantages predicted for asexual lineages. Unlike other asexual animal taxa, genomes of oribatid mites show reduced accumulation of slightly deleterious mutations compared to their sexual relatives, possibly facilitated by large population sizes (Maraun *et al*., 2012;Jaron *et al*., 2020b; Brandt *et al*., 2017; Bast *et al*., 2018). Also, similar to other asexual organisms, oribatid mites show no increased abundance and activity of transposable elements compared to sexuals (Bast *et al*., 2016, 2019). These findings suggest that asexual oribatid mites indeed escape the dead-end fate usually associated with asexual lineages.

## Methods

### Animal sampling and DNA/RNA extraction

Animals were sampled in the fall of 2015 and 2017 from leaf litter and soil at four different forest sites in Germany (Goettingen forest (GF), Hainich (H), Kranichstein forest (KF), Schwaebische Alb (SA); for details see **Supplementary Table S6**). Living animals were separated from the leaf litter with heat gradient extraction (Kempson *et al*., 1963) and identified (Weigmann, 2006), followed by at least one week of starving to reduce potential contaminants derived from gut contents. Afterwards, animals were cleaned by removing surface particles in sterile water, several minutes of washing in a solution of hexane:bleach:detergent:water (25:25:1:49) and rinsing with sterile water before extraction. Note that animals were alive after cleaning.

For generating reference genomes, DNA was extracted from one single individual of *O. subpectinata* collected in 2015 from GF leaf litter and *O. nova* collected in 2015 from KF leaf litter using the QIAamp DNA Micro kit according to manufacturer’s instructions. To generate transcriptomes for annotation of reference genomes, RNA was extracted from five pooled individuals per species from the same collection batch. For this, individuals were frozen in liquid nitrogen and, after addition of Trizol (Life Technologies), mechanically crushed with beads (Sigmund Lindner). Next, chloroform and ethanol was added to the homogenised tissue and the aqueous layer transferred to RNeasy MinElute Columns (Qiagen). Subsequent steps of RNA extraction were done following the RNeasy Mini Kit protocol, including DNAse digestion. Finally, RNA was eluted into water and stored at −80°C. To infer haplotype divergence, RNA was extracted from single individuals of *O. nova* and *O. subpectinata* from H, SA and KF (three individuals from each forest site for each species) from the 2017 collection batch. RNA extraction was done as described above. DNA and RNA quantity and quality was measured using respectively Qubit and NanoDrop (Thermo Scientific), and Bioanalyzer (Agilent).

### Reference genome assemblies and contaminant removal

For genome sequencing, extracted DNA from single individuals was amplified in two independent reactions using the SYNGIS TruePrime WGA kit and then pooled. Four libraries were generated for each reference genome (three paired end libraries with average insert sizes of respectively 180, 350 and 550bp, and a mate-pair library with 3000 bp insert size). Libraries were prepared using the illumina TruSeq DNA or Nextera Mate Pair Library Prep Kits, following manufacturer instructions, and sequenced on the Illumina HiSeq 2500 system, using v4 chemistry and 2x 125 bp reads at FASTERIS SA, Plan-les-Ouates, Switzerland. This resulted in a total number of 451*10^6^ reads for *O. nova* with a total read coverage of 490-fold and 387*10^6^ reads for *O. subpectinata* with a total read coverage of 420-fold (for details see **Supplementary Table S2**). Read quality trimming and adapter clipping of paired reads was done using Trimmomatic v0.36 (Bolger *et al*., 2014) with the following options: ILLUMINACLIP:/all-PE.fa:2:30:10 LEADING:20 TRAILING:20 SLIDINGWINDOW:3:20 MINLEN:100. This resulted in 56% and 46% surviving read pairs (for details see **Supplementary Table S2**). For mate pair quality trimming, Nxtrim v0.4.1 (O’Connell *et al*., 2015) with options --separate --preserve-mp --minlength 40, followed by Trimmomatic v0.36 with options ILLUMINACLIP:/all-PE.fa:2:30:10 LEADING:20 TRAILING:20 SLIDINGWINDOW:4:20 MINLEN:60 were used to identify properly paired reads and to remove low quality bases and adapters. This resulted in 54% and 48% surviving read mate pairs (for details see **Supplementary Table S2**).

With the available read data, we tested a range of assembly strategies. The best assemblies were generated using normalized overlapped reads, because whole genome amplification introduces overrepresented genomic regions, which leads to coverage bias that is problematic for assembly. Overlapped read libraries were generated by merging the paired forward and reverse reads of the 180 bp read libraries and additionally merging unpaired reads, followed by normalization using BBnorm v37.82 (Bushnell, 2014). These normalized overlap read libraries were assembled into contigs using SPAdes v3.10.1, a multi k-mer assembler (Bankevich *et al*., 2012), with options -m 400 --careful -k 21, 33, 55, 77, 99, 111, 127. The resulting contigs were ordered into scaffolds using the 350 bp, 500 bp and 3000 bp read libraries using SSPACE v3.0 (Boetzer *et al*., 2011) with default parameters. To close gaps emerging during scaffolding, GapCloser v1.12 (Luo *et al*., 2012) with option -l 125 was run. For details see https://github.com/AsexGenomeEvol/HD_Oppiella: assembly and mites.

Scaffolds that were likely from contaminants (e.g., bacteria, fungi) were removed by first annotating and visualizing contaminations using BlobTools v1.0 (Laetsch & Blaxter, 2017), followed by custom filtering. For this, coverage of each scaffold was estimated by mapping reads back to the scaffolds using bwa mem v0.7.15 (Li, 2013) and coverage calculated with BBTools v73.82 (Bushnell, 2014). Additionally, for annotation, scaffolds were blasted using ncbi-blast v2.7.1+ blastn with options -outfmt ‘6 qseqid staxids bitscore evalue std sscinames sskingdoms stitle’ -max_target_seqs 10 -max_hsps 1 -evalue 1e-25 against the nt database v 2016-06. Scaffolds without hits to metazoans were filtered out from the assemblies using a custom script (see https://github.com/AsexGenomeEvol/HD_Oppiella: contamination_filtration.py). Next, scaffolds were sorted by decreasing length, scaffold headers renamed and scaffolds shorter than 500 bp removed, resulting in the final assemblies (v03). The assemblies were checked for quality and completeness by calculating standard genome statistics and by checking presence, fragmentation and duplication of arthropod core genes using CEGMA v2.5 and BUSCO v3.0.2 (Parra *et al*., 2007; Simão *et al*., 2015; Seppey *et al*., 2019). For details see **Supplementary Table S1**.

### Genome annotation

The de-contaminated genome assemblies v03 were annotated using MAKER v2.31.8 (Holt & Yandell, 2011), a hybrid *de novo* evolution-based and transcript-based method. For this, repetitive genomic regions are first masked using RepeatMasker v4.0.7 (Smit *et al*., 2013-2015) as implemented in MAKER. Protein coding genes were then predicted in a 2-iterative way described in (Campbell *et al*., 2014) with minor modifications following author recommendations. For the first iteration, genes were predicted using Augustus v3.2.3 (Stanke *et al*., 2006) trained with the BUSCO v3.0.2 results (arthropoda_odb9 lineage with the --long option). A combination of UniProtKB/Swiss-Prot (release 2018_01) and the BUSCO arthropoda_odb9 proteome were used as protein evidence. The Trinity assembled mRNA-seq sequences (described below) were used as transcript evidence. The resulting gene models from iteration 1 were then used to retrain Augustus as well as SNAP v2013.11.29 (Korf, 2004) and a second iteration was performed. Following, predicted protein coding genes were functionally annotated using Blast2GO v5.5.1 (Conesa *et al*., 2005) with default parameters against the NCBI *non-redundant arthropods* protein database and the *Drosophila melanogaster* (drosoph) database v 2018-10. The MAKER configuration files are available at https://github.com/AsexGenomeEvol/HD_Oppiella.

For the MAKER annotation, RNAseq reads were quality trimmed with Trimmomatic v0.36 with options adapters.fa:2:30:12:1:true LEADING:3 TRAILING:3 MAXINFO:40:0.4 MINLEN:80. For generating genome-guided transcriptome assemblies, trimmed reads were first mapped against the genomes using STAR v2.5.3a (Dobin *et al*., 2013) under the ‘2-pass mapping’ mode and default parameters. Following, the outputs were used with Trinity v2.5.1 (Haas *et al*., 2013) set to ‘genome guided’ mode (parameters: --genome_guided_max_intron 100000 --SS_lib_type RF --jaccard_clip). For quality filtering of the resulting transcriptomes, the trimmed RNAseq reads were mapped back against the transcriptomes using Kallisto v0.43.1 (Bray *et al*., 2016) with options --bias and --rf-stranded, then transcripts with at least 1 TPM in any samples were retained. All computation for genome assembly and annotation were run on the Vital-IT cluster of the Swiss Institute of Bioinformatics.

### Haplotype divergence: RNAseq, quality control and mapping

RNA extracts were fragmented to 175 nt for strand-specific library preparation using the NEBNext^®^ Ultra^™^II Directional RNA Library Prep Kit. Paired-end sequencing with a read length of 100 bp was performed on a HiSeq2000 platform at the GTF (Genomics Technology Facility Lausanne, Switzerland). Data processing for haplotype divergence inference was done using the high performance computing cluster of the Gesellschaft für Wissenschaftliche Datenverarbeitung Goettingen, Germany (GWDG). RNAseq reads were adapter- and quality-trimmed using TrimGalore v0.6.5 with default options (Phred quality threshold 20; adapter auto-detection; Martin, 2011; Krueger, 2012). Contaminating sequences were removed using kraken2 (--paired; --db minikraken2_v2; Wood & Salzberg, 2014) followed by mapping paired-end reads of each individual simultaneously against their respective reference genome, scaffolds flagged as contaminating sequences assembled together with the respective mite reference genomes (identified as described above), the human reference genome GRCh38.p12 (GenBank assembly accession: GCA_000001405.27) and the human microbiome (downloaded from https://www.hmpdacc.org/hmp/HMREFG/all/index.php) using bbmap v37.66 (bbsplit; maxindel=100k; ambiguous=best; Bushnell, 2014). Portions of reads were found to be derived from contaminating RNA of human and microbial origin with fractions ranging from 40.36% to 90.31% in *O. subpectinata* and 53.33% to 93.04% in *O. nova* (see **Supplementary Table S7**). Only oribatid-mite-exclusive reads, i.e., read pairs that mapped best and unambiguously against the mite reference genomes were kept for further processing and mapped to the reference genomes using STAR v2.7.3a with standard parameters. All commands are available under https://github.com/AsexGenomeEvol/HD_Oppiella: mapping.U.

### Haplotype divergence: Variant calling

Read-group information was added and PCR and optical duplicates were removed from mapped reads using Picard v2.20.2 (Broad Institute, 2020). Reads without a mapping mate were deleted using samtools view (Li *et al*., 2009) and reads sorted by coordinate using GATK v4.0.3.0 SortSam (McKenna *et al*., 2010). Next, the nine thus filtered alignments per species were merged with samtools merge for subsequent SNP calling. Sequences spanning intronic regions were removed using GATK SplitNCigarReads. GATK HaplotypeCaller was run per individual with -ERC set to BP_RESOLUTION to enable calling of non-variant sites and --dont-use-soft-clipped-bases to exclude soft-clipped overhangs from SplitNCigarReads. Individual gvcf-files were combined into one species-gvcf-file using GATK CombineGVCFs. Genotypes were called using GATK GenotypeGVCFs and the option -allSites. All commands used are available under https://github.com/AsexGenomeEvol/HD_Oppiella: calling.U

### MDS plots and AMOVA

For calculating MDS plots and AMOVA, first genotypes with a coverage < 10 were removed from gvcf-files using vcftools v0.1.15 (Danecek *et al*., 2011). Next, sites including at least one missing genotype, monomorphic or tri-allelic variants, and indels were removed. To visualise genotype composition of populations, multidimensional scaling (MDS; two scales for two-dimensional representation) was done using plink v1.9 with the options --cluster, --mds-plot 2 eigvals and --allow-extra-chr (Purcell *et al*., 2007). Population differentiation was tested based on the filtered set of SNPs with an Analysis of Molecular Variance (AMOVA) and a randomness test with the packages vcfR and poppr in R (R Core Team, 2013; Kamvar *et al*., 2014; Mateus & Caeiro, 2014; Knaus & Gruenwald, 2017). All commands used are available under https://github.com/AsexGenomeEvol/HD_Oppiella: MDS.U, AMOVA.R.

### Heterozygosity, Fis and SFS

For calculating the percentage of heterozygous genotypes per individual, variants were filtered as described above (except monomorphic variants were not excluded). The percentage of heterozygous positions per individual was calculated using unix commands. Similarly for F_is_ and SFS calculation, first genotypes with a coverage < 10 were removed from gvcf files using vcftools v0.1.15. For F_is_ calculation the gvcf-file was next subset into lineages I and II for *O. nova* and populations for *O. subpectinata* using vcf-subset (vcftools). For SFS calculation, subsetting was done for seven individuals of *O. nova* potentially showing the Meselson effect (see **Results**) with F_is_ < 0. Afterwards, sites including at least one missing genotype, monomorphic or tri-allelic variants, and indels were removed from the different subsets. F_is_ was calculated based on the filtered sets of SNPs using vcftools (option --het). The SFS was calculated using Pop-Con and standard parameters (Anselmetti et al *in prep* https://github.com/YoannAnselmetti/Pop-Con). The fold excesses of shared heterozygous

SNPs over HWE were estimated by running Pop-Con with the option -fold for a range of parameters and comparing the specific genotype profiles for an indication of excess (shown as part of expected SFS). All commands used are available under https://github.com/AsexGenomeEvol/HD_Oppiella: heterozygosity.U, Fis.U and SFS.U.

### Haplotype phasing

For phasing, variants were filtered as described above (except only sites completely missing any genotype information, i.e. all individuals missing a genotype, were excluded). Phasing of haplotypes was done per individual using phASER v1.1.1 with minimum mapping quality of reads set to 30, minimum base quality set to 20 and bottom cut-off to quantile for alignment score set to 0 in paired-end mode for each individual, separately, utilising heterozygous variants with minimum coverage of 10 for each individual (Castel *et al*., 2016). Haplotypes with < 10 unique reads mapping were removed from the output data. Output data from phASER were converted into haplotype sequences for the corresponding positions in the reference genome using a custom script (available under https://github.com/AsexGenomeEvol/HD_Oppiella: meselson.py), which extracts the corresponding sequence from the reference genome and generates the two haplotypes by modifying the extracted sequence with the haplotype information from phASER. Furthermore, as some SNPs might not have been called by GATK HaplotypeCaller due to insufficient coverage, all bases of the reference genome with coverage < 10 - the coverage threshold for genotype filtration - were excluded from further analysis (changed to N; coverage estimated with bedtools version 2.26.0 from individual bam-files after removing overhanging N’s at read ends using GATK SplitNCigarReads with the option --process-secondary-alignments). All regions comprising at least one phase per individual that overlapped with a phase of a different individual by at least 100 bp were included for downstream analyses (forming continuous stretches of phases; see **Results**). Haplotypes were labeled according to their divergence from the reference genome (haplotype A being closer to the reference genome, haplotype B being more diverged). For this, positions including degenerate bases were deleted using trimAl (Capella-Gutiérrez *et al*., 2009) and the pairwise distance of each haplotype to the reference genome was calculated using snp-dist (Seeman, 2020). Only thus modified phaseable regions >= 100bp were included for downstream analyses. All commands and two scripts used are available under https://github.com/AsexGenomeEvol/HD_Oppiella: phasing.U, refgenomedist.U, extract.pl and convert_fasta.py.

To identify putative paralogs in the phased regions, these regions were tested for double coverage compared to the genomic baseline. Reads used to assemble genomes were mapped back to single copy genes and duplicated genes identified by BUSCO (see above), and additionally to the phased regions and to scaffolds from which the phased regions were derived (but that were masked in the phased regions), using bowtie2 v2.3.4.1 with standard parameters (Langmead & Salzberg, 2012). The mapped alignments were quality filtered (MAPQ score > 10) using samtools and optical and PCR duplicates removed using Picard Tools Mark Duplicates v2.22.0. Following, coverage was calculated using bedtools genomecov v2.26.0 (Quinlan & Hall, 2010).

### Topology testing

To enable testing whether alignments of phaseable regions are better explained by a topology separating the haplotypes (as expected under asexuality) as compared to a topology separating populations (expected under sexual reproduction) or *vice versa*, a constrained tree search was done. Two constrained Maximum Likelihood (ML) trees, one complying with a fixed haplotype-separating topology (asex-tree), the other with a fixed population-separating topology (sex-tree) were calculated for each phased region using iqtree v1.6.10 with 1000 bootstrap replicates and model-testing included (Nguyen *et al*., 2015). For *O. nova*, we restricted the analysis to the seven individuals representing the two divergent lineages (see Results; **Figure 2a**). The *O. nova* asex-trees were constrained to separate the haplotypes A & B at its base, lineages I & II per haplotype and finally the populations per lineage and haplotype (for the constraining tree, see https://github.com/AsexGenomeEvol/HD_Oppiella: Onasex.tre). The sex-trees were constrained to separate the lineages I & II at its base and the populations per lineage (no haplotype separation; for the constraining tree, see https://github.com/AsexGenomeEvol/HD_Oppiella: Onsex.tre). For *O. subpectinata* the asex-trees were constrained to separate the haplotypes A & B at its base and the populations per haplotype (to provide an unrooted tree including a trichotomy the haplotypes B of the most divergent population SA were separated from haplotypes B of the other two populations; for the constraining tree, see https://github.com/AsexGenomeEvol/HD_Oppiella: Osasex.tre). The sex-trees were constrained to separate exclusively the populations (no haplotype separation; for the constraining tree, see https://github.com/AsexGenomeEvol/HD_Oppiella: Ossex.tre). Next, the distance between the two resulting trees and an unrestricted best fitting ML tree was estimated according to Kuhner & Felsenstein, 1994 with the dist.topo function implemented in the R package ape (Paradis & Schliep, 2019). To enable the comparison between distances of different phaseable regions, the topological distances of the best fitting ML tree to the haplotype-separating tree and to the population-separating tree were combined as Δ _dist. HD-tree - dist. sex-tree_ value for each phaseable region. To test if the haplotype-separating tree was a significantly better fit to the alignment than the population-separating tree, trees were compared using RELL approximation with 10,000 RELL replicates and an approximately unbiased (AU) test with iqtree (Kishino *et al*., 1990; Shimodaira, 2002). Using the AU test, we also compared both constrained trees to the unconstrained tree. For detailed information, see https://github.com/AsexGenomeEvol/HD_Oppiella: treecalcandtopotest.U and topodist.UR.

### Parallel divergence testing

Phasing and haplotype reconstruction was done as described above but coverage was reduced to a minimum of five to increase the number of informative sites, and thereby the number of non-polytomous trees. Calculation of best fitting ML trees was done as described above. Resulting topologies were screened by eye for being non-polytomous, for showing haplotype separation and for parallel divergence. To calculate the probability to observe parallel divergence in at least eight trees out of 31 haplotype separating eight taxa trees by chance (lineage I) we used the binomial theorem 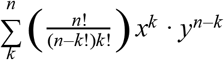 with n = 31 (31 trees showing haplotype separation), k = 8 (8 trees showing parallel divergence), x = 0.067 (the probability to observe parallel divergence in two rooted four taxa trees by chance) and y = 0.933 (counter event; 1-x). To calculate the probability to observe parallel divergence in at least 38 trees out of 55 haplotype separating six taxa trees by chance (lineage II) we used the binomial theorem with n = 55, k = 38, x = 0.333 and y = 0.667.

### Statistical analyses

All statistical analyses were done in R v3.6.3 (R Core Team, 2013) unless mentioned otherwise.

## Supplementary Information

### Supplementary Figures

**Supplementary Figure 1:**
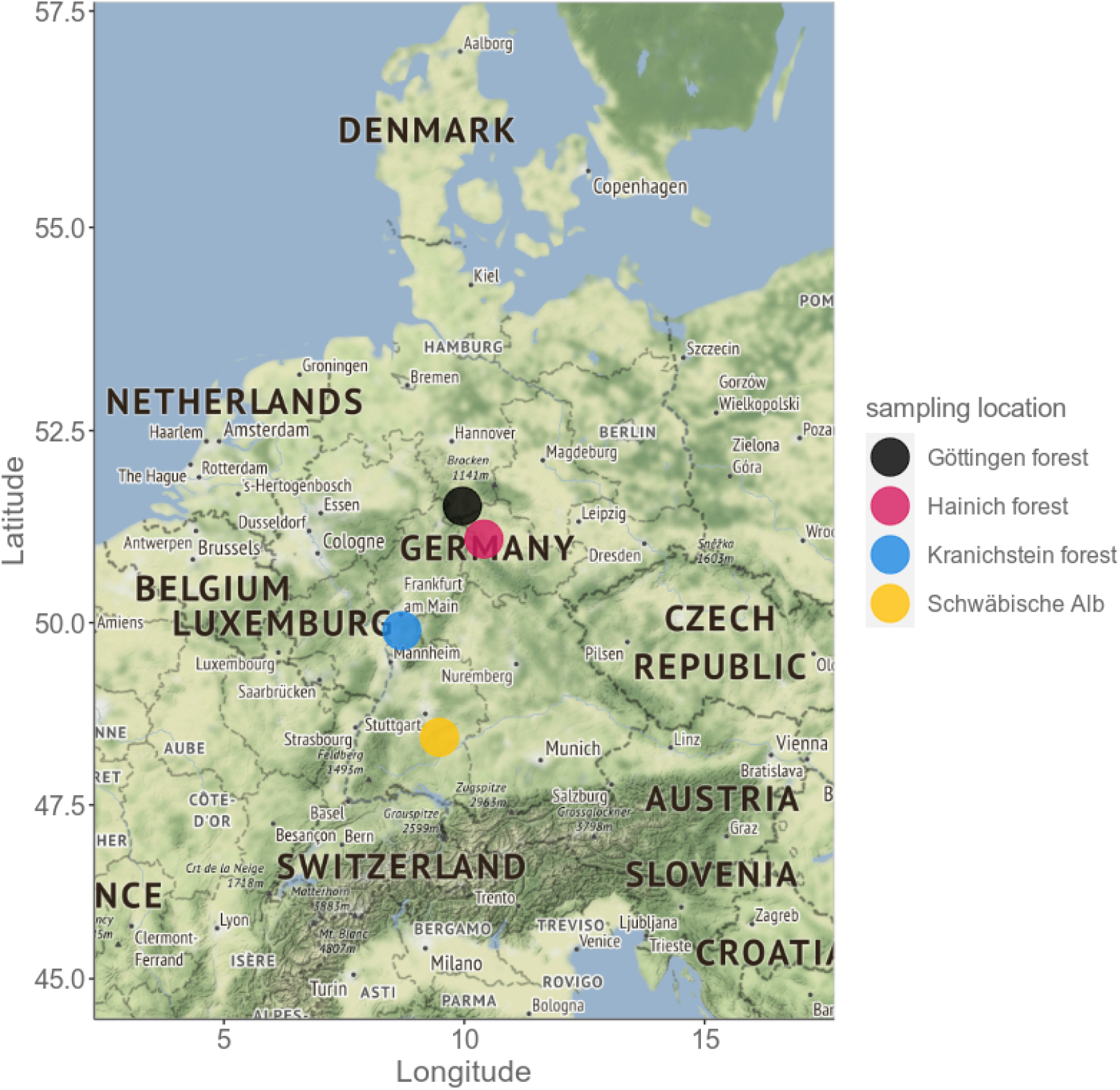
The oribatid mite samples were collected in different forests in Germany. For detailed information on sampling, see Supplementary Table S8.

**Supplementary Figure 2:**
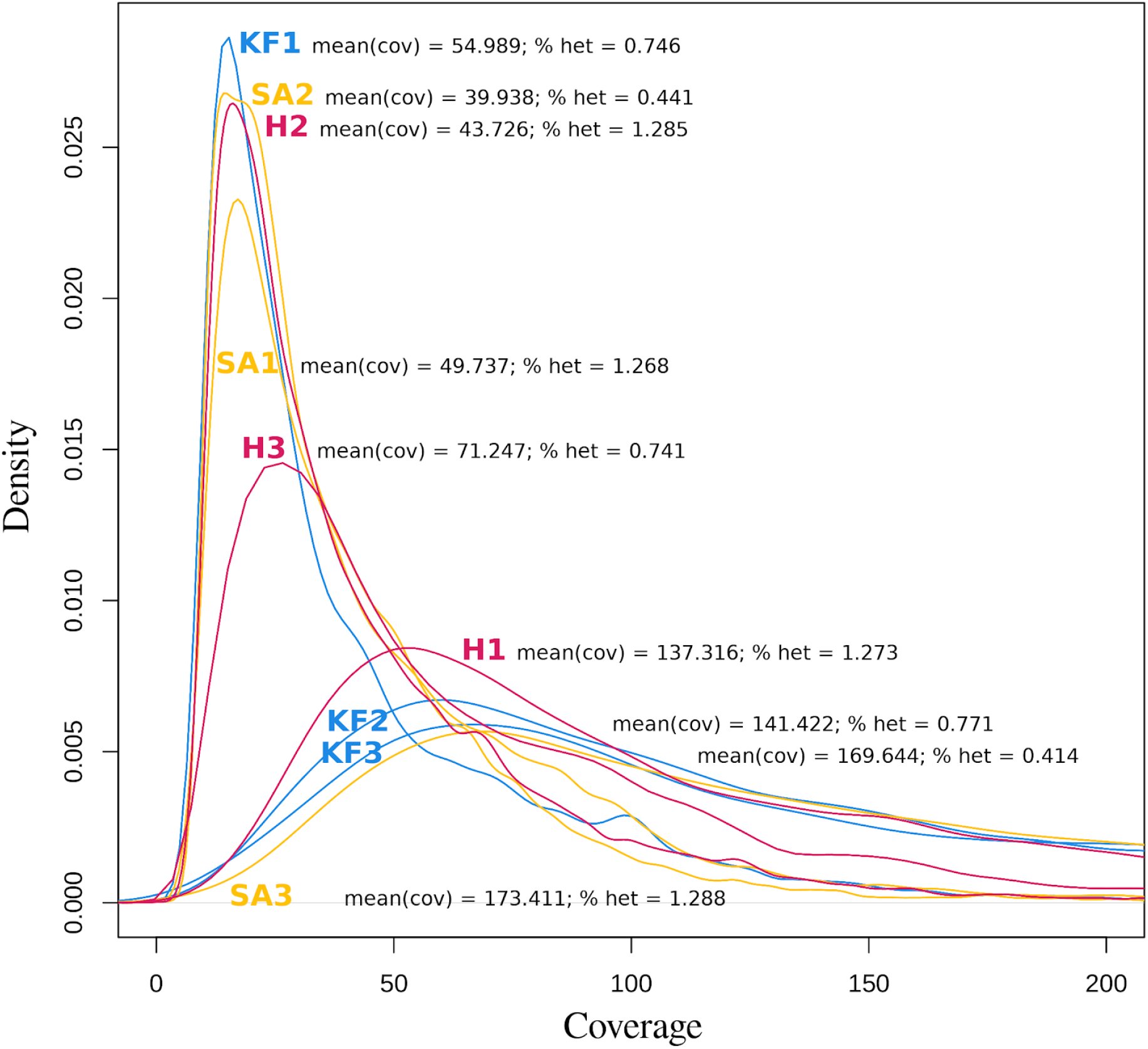
Kernel density of genotype coverage per individual. There is no significant correlation between percentages of heterozygous sites (% het) and mean coverage of genotypes (mean(cov)) per individual (linear model *F* = 0.00144; *P* = 0.971) indicating that the large differences in heterozygosity between individuals are not driven by coverage variation. H Hainich; KF Kranichstein forest; SA Schwaebische Alb.

**Supplementary Figure 3:**
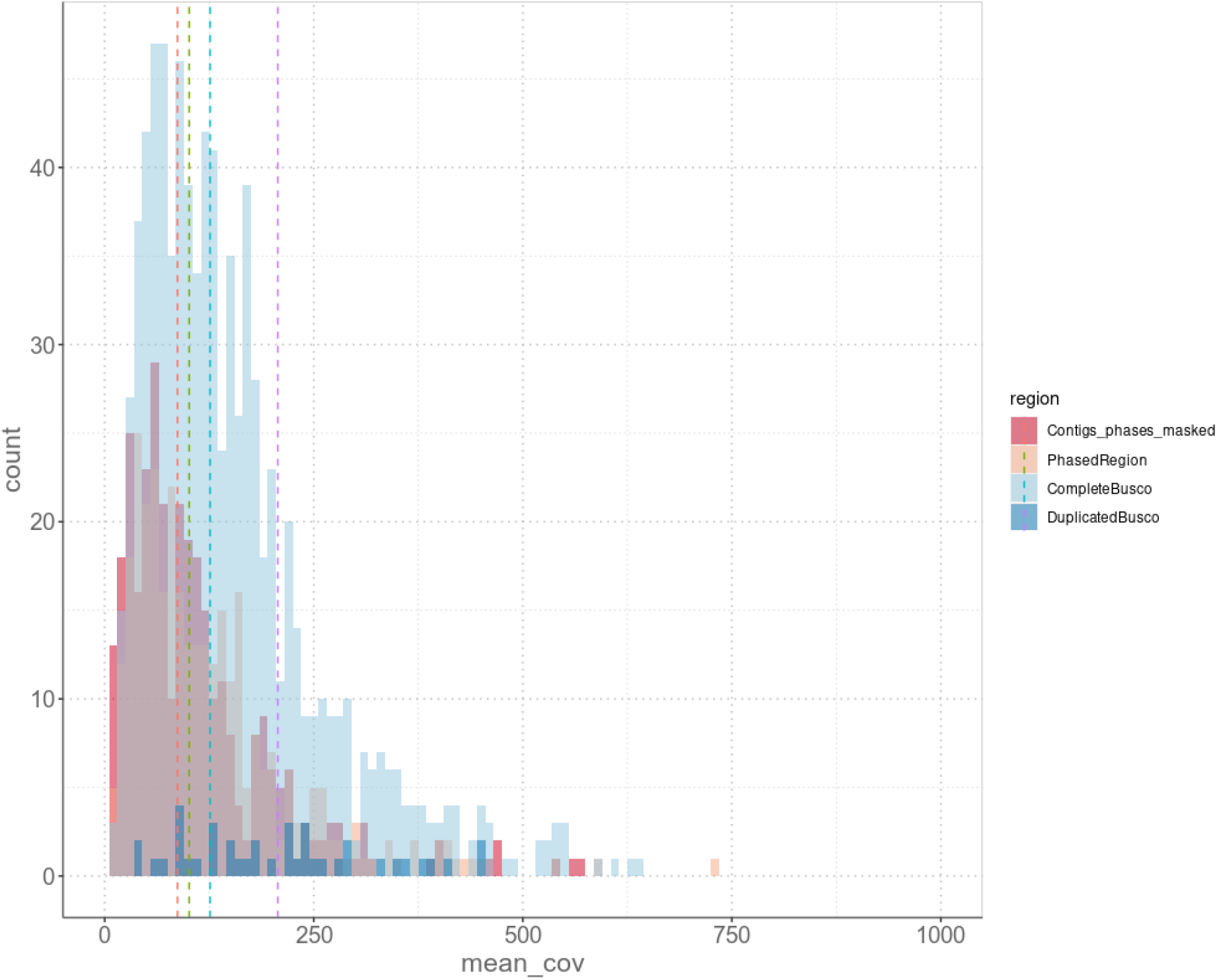
Coverage distributions indicate that phased regions are not enriched for merged paralogs. Count frequencies of coverages of the different regions based on genomic read data: Contigs that contained the phased regions but were masked for these, the phased regions, complete single-copy genes and duplicated genes as identified by BUSCO. Dotted lines represent the median of the coverage for the respective regions.

### Supplementary Tables

**Supplementary Table S1:** Genome statistics and completeness scores. v02: decontaminated assembly, v03: decontaminated assembly with > 500bp scaffold length size filter

| Species | CEGMA | Busco | Genome Size [Mbp] | Scaffold N50 [bp] | Number of scaffolds | Largest scaffold length [bp] | GC content [%] | N [%] |
| --- | --- | --- | --- | --- | --- | --- | --- | --- |
| *O. nova* |  |  | 207.08 | 6604 | 83865 |  |  | 1.64 |
| Blobtools v02 | C: 88.31<br>P: 95.56 | C:87.1%[S:78.8%,<br>D:8.3%],F:6.6%,<br>M:6.3%,n:1066 | 202.44 | 6493 | 82784 | 406770 | 30.41 | 1.68 |
| Filtered v03 | C: 88.31<br>P: 95.56 | C:87.5%[S:78.9%,<br>D:8.6%],F:6.6%,<br>M:5.9%,n:1066 | 196.72 | 6753 | 63118 | 406770 | 30.37 | 1.73 |
| *O. subpectinata* |  |  | 217.82 | 6873 | 73173 |  |  | 1.44 |
| Blobtools v02 | C: 88.31<br>P: 97.58 | C:86.6%[S:78.6%,<br>D:8.0%],F:6.3%,<br>M:7.1%,n:1066 | 217.02 | 6852 | 73101 | 165769 | 30.84 | 1.44 |
| Filtered v03 | C: 88.31<br>P: 97.58 | C:86.2%[S:78.3%,<br>D:7.9%],F:6.4%,<br>M:7.4%,n:1066 | 213.17 | 7017 | 60250 | 165769 | 30.83 | 1.47 |

**Supplementary Table S2:** Information on the read library insert sizes, number of reads generated, estimated coverage, and number of surviving read pairs after filtering.

| Species | Library | No of raw read pairs | Coverage | No of read pairs surviving |
| --- | --- | --- | --- | --- |
| *O. nova* | 180 | 125889255 | 137 | 95460017 (75.83%) |
|  | 350 | 73798941 | 80 | 44494871 (60.29%) |
|  | 550 | 76545499 | 83 | 34976565 (45.69%) |
|  | 3000 | 174946393 | 190 | 33642006 (53.99%) |
| *O. subpectinata* | 180 | 112721137 | 122 | 89023834 (78.98%) |
|  | 350 | 87673896 | 95 | 65281720 (74.46%) |
|  | 550 | 71743398 | 78 | 40465571 (56.40%) |
|  | 3000 | 115207485 | 125 | 21898456 (53.96%) |

**Supplementary Table S3:** Annotation statistics

| Species | Gene number | mRNA number | Gene min length [bp] | Gene mean length [bp] | Gene max length [bp] | Genome covered by genes [%] | Exon mean length [bp] | Intron mean length [bp] |
| --- | --- | --- | --- | --- | --- | --- | --- | --- |
| *O. nova* | 23761 | 24130 | 30 | 1956 | 42876 | 23.6 | 258 | 270 |
| *O. subpectinata* | 23555 | 23803 | 49 | 2148 | 53142 | 23.7 | 258 | 246 |

**Supplementary Table S4:** Detailed information on successfully phased regions. **\*** No. phased regions after removal of sites with coverage < 10; **#** No. phased regions after removal of regions with sequence similarity too large for tree calculation; **∼** No. significantly haplotype divergence positive regions.

| Species | No. phased regions | total length | * | total length | median length | # | total length | ~ | total length |
| --- | --- | --- | --- | --- | --- | --- | --- | --- | --- |
| *O. nova* | 329 | 1,233,469 | 281 | 140,966 | 358 | 223 | 127,123 | 69 | 37,693 |
| *O. subpectinata* | 302 | 2,010,902 | 275 | 206,255 | 563 | 268 | 204,851 | 1 | 115 |

**Supplementary Table S5:** Results of ML based tree topology (AU) tests. The table lists numbers of haplotype sequence alignments of phased regions that provide a significantly better fit to a constrained tree separating haplotypes than to a constrained tree separating populations (asex-tree + sex-tree -) and vice versa (asex-tree - sex-tree +) for the asexual species *O. nova* and its sexual relative *O. subpectinata*. To further assess how consistent the topology of the best ML tree was with either one of the constrained trees, we additionally ran AU tests using the unconstrained tree (e.g. unconstrained tree: asex-tree + sex-tree - indicates non-significant differences between the unconstrained tree and the asex-tree). For some phased regions AU tests could not be run due to insufficient variation between haplotypes.

|  | *O. nova* | *O. subpectinata* |
| --- | --- | --- |
| asex-tree + sex-tree - | 69 | 1 |
| asex-tree - sex-tree + | 70 | 258 |
| both + (non-significant) | 70 | 4 |
| both - (test impossible) | 14 | 5 |
| unconst.: asex-tree + sex-tree - | 12 | 0 |
| unconst.: asex-tree - sex-tree + | 10 | 95 |

**Supplementary Table S6:**
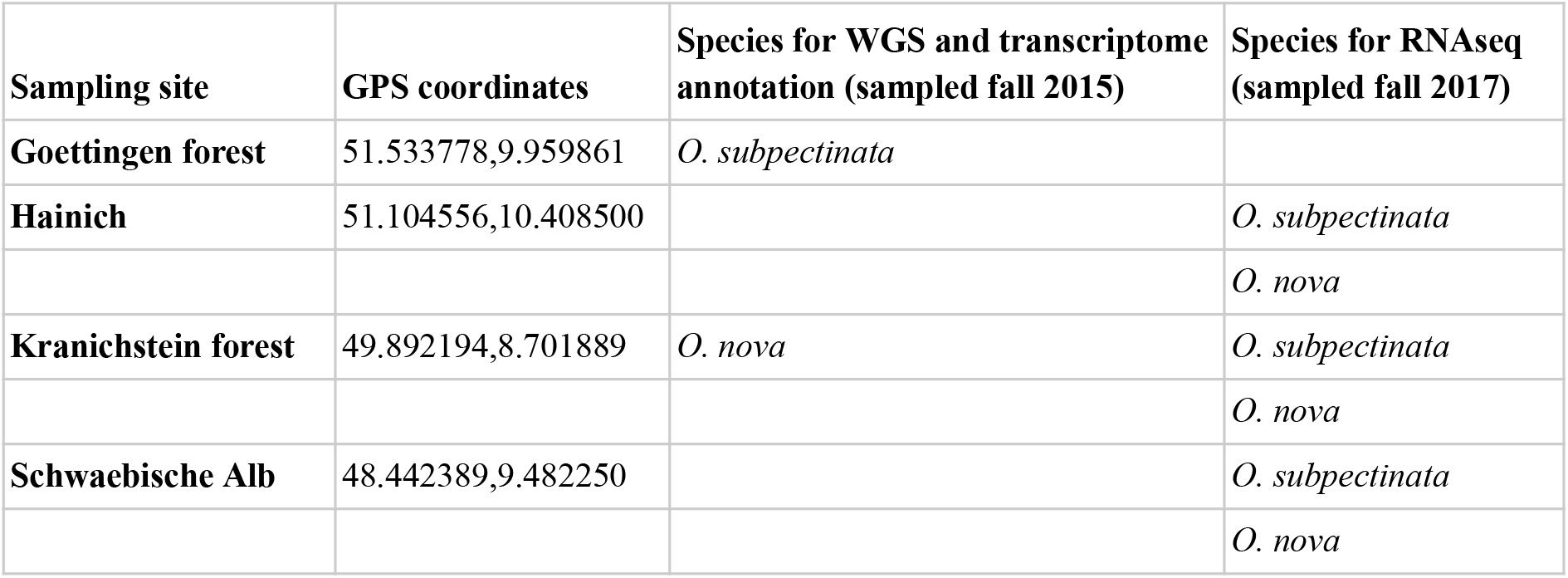
Additional information on sampling sites and species.

**Supplementary Table S7:** numbers of read pairs/reads after different steps of quality trimming, contamination removal and mapping.

| Species | Population | Individual | No. raw read pairs | No. read pairs after trimming | No. read pairs after contaminant removal | % contaminating read pairs | No.mapped reads after duplicate removal |
| --- | --- | --- | --- | --- | --- | --- | --- |
| *O. nova* | Hainich | H1 | 12,305,768 | 10,922,239 | 5,073,080 | 53.33 | 1,538,502 |
|  |  | H2 | 11,653,023 | 10,150,566 | 1,546,946 | 84.76 | 434,144 |
|  |  | H3 | 17,293,884 | 14,456,993 | 3,959,499 | 72.61 | 866,110 |
|  | Kranichstein forest | KF1 | 14,818,283 | 12,790,718 | 1,426,174 | 88.85 | 444,750 |
|  |  | KF2 | 14,514,107 | 12,380,574 | 4,874,135 | 60.63 | 1,780,426 |
|  |  | KF3 | 22,424,703 | 20,094,950 | 7,214,778 | 64.1 | 2,138,804 |
|  | Schwaebische Alb | SA1 | 10,460,975 | 9,570,865 | 1,339,805 | 86 | 567,076 |
|  |  | SA2 | 21,489,919 | 19,714,647 | 1,371,202 | 93.04 | 500,764 |
|  |  | SA3 | 23,061,245 | 21,030,385 | 7,930,318 | 62.29 | 2,083,292 |
| *O. subpectinata* | Hainich | H1 | 21,756,064 | 20,776,464 | 5,399,072 | 74.01 | 2,828,074 |
|  |  | H2 | 18,105,719 | 15,780,873 | 5,727,395 | 63.71 | 1,320,928 |
|  |  | H3 | 12,584,357 | 11,011,539 | 2,464,755 | 77.62 | 997,412 |
|  | Kranichstein forest | KF1 | 26,438,129 | 23,518,688 | 4,829,639 | 79.46 | 3,472,828 |
|  |  | KF2 | 36,588,456 | 34,945,277 | 3,386,946 | 90.31 | 4,238,092 |
|  |  | KF3 | 28,975,922 | 23,341,626 | 11,877,834 | 49.11 | 3,279,550 |
|  | Schwaebische Alb | SA1 | 21,616,073 | 20,271,910 | 12,090,364 | 40.36 | 4,230,422 |
|  |  | SA2 | 12,186,958 | 10,229,333 | 2,695,017 | 73.65 | 739,384 |
|  |  | SA3 | 15,924,247 | 13,972,229 | 3,810,147 | 72.73 | 1,178,238 |

## Code availability

All scripts and commands as referred to in the article are available at https://github.com/AsexGenomeEvol/HD_Oppiella.

## Data availability

Reference genomes and annotations of the two species are available in the European Nucleotide Archive (ENA; accession numbers PRJEB39968). Transcriptomes and RNAseq reads of the two species used for inferring allelic sequence divergence and for annotations are available from SRA (accession numbers PRJNA662767).

## Acknowledgements

We like to thank Juergen Brandt for adjusting figures to the needs of people with colour vision deficiency and Christoph Bleidorn for advice regarding topology testing. This study was supported by core funding of S.S., by DFG Emmy Noether fellowship BA 5800/3-1 to J.B., Swiss SNF grant CRSII3_160723 to T.S., M.R.R. and N.G., and Swiss SNF grant PP00P3_170627 to T.S.

## Contributions

A.B., J.B. and T.S. conceived and designed the study. A.B. and C.B. collected samples and determined species. J.B., Z.D. and M.L. performed wet lab work. A.B., J.B. and P.TV. performed data analysis with input from T.S. and S.S. T.S., N.G., M.R.R., K.S.J., C.M.F., M.M., B.H., E.F., D.J.P., I.S., Y.A., P.S.and S.S. contributed to data interpretation and analyses and A.B., J.B. and T.S. wrote the paper with input from all authors.

